# Spatial transcriptomics reveals interferon pathway enrichment in early relapsing serous carcinoma

**DOI:** 10.64898/2026.10.12.758400

**Authors:** Franziska Niemeyer, Niina Aho, Satu Hänninen, Omar Youssef, Olli Mikael Carpén, Andrew Erickson, Anna Ray Laury

## Abstract

High-grade serous carcinoma (HGSC) is characterized by extensive inter and intra-patient heterogeneity, complicating the identification of reliable prognostic biomarkers and the prediction of treatment response. In clinical practice, the origin of diagnostic biopsies varies, from tubo-ovarian (adnexal) tissue to omental or peritoneal metastatic foci. Whether the different sites provide similar information for molecular characterization or treatment decisions remains unclear. Studies comparing site-specific molecular details and the spatial relationship of HGSC tissue from patients with different treatment outcomes could provide answers to the open questions. Here, we used spatial transcriptomics to profile matched treatment-naïve adnexal and omental tumor samples from nine HGSC patients stratified by platinum-free interval (PFI). Pathway enrichment analysis revealed several differences between short and medium- and long PFI tumors, however most of these differences were site-agnostic. Most strikingly, tumor epithelium from short PFI patients showed an inflammatory phenotype driven by increased interferon response signaling, carried by effector interferon-stimulated gene expression at both tumor sites. Further, medium- and long PFI tumors were enriched for proliferative and cell cycle regulatory programs. Differential gene expression and copy-number analyses identified three genes, complement gene C3, and interferon-related genes IFI27 and BST2, consistently associated with short PFI across adnexal and omental sites. These were validated at the tissue level using RNA in-situ hybridization. A case study of copy-number-defined subclones further suggested that this phenotype can arise as a distinct subclonal population within a single tumor. Outcome analysis, using two publicly available ovarian cancer cohorts, showed that high C3 expression was significantly associated with shorter platinum-free survival. Together, these findings implicate complement and interferon signaling as concurrent features of treatment-naïve HGSC tumors with early relapse in both adnexal and omental tissue, and nominate C3 as a candidate prognostic biomarker and potential therapeutic target.

## Introduction

Ovarian cancer is the 8th most frequent cancer in women globally, with about 331,000 new cases diagnosed annually^1^. High-grade serous carcinoma of the fallopian tube and ovary (HGSC) is the most common and most lethal ovarian cancer subtype, accounting for 65-70% of ovarian cancers and approximately 80% of ovarian cancer deaths^2,3^. Patients are often diagnosed at advanced stage, when the cancer has already spread to the abdomen^4^. Most patients initially respond to chemotherapy, however, this is nearly always followed by eventual relapse and rapid disease progression. Determining who will relapse, and when, is not currently possible using routine clinicopathological information, and molecular assays are not in clinical use for this purpose, apart from homologous recombination deficiency (HRD)-testing as a genomic assay for treatment stratification. The inability to predict treatment response presents a significant clinical challenge, highlighting the need for reliable prognostic biomarkers^5,6^.

Early comparisons of matched pelvic and abdominal tumors reported only subtle differences in gene expression from bulk and single-cell RNA-sequencing data^7,8^, but larger multi-region studies have since shown that the immune and inflammatory phenotype of a tumor depends partly on its anatomic site: compositional, topological, and functional differences all distinguish adnexal tumors from distant peritoneal disease^9^. Recent studies comparing transcriptomic data in the context of overall survival have found significant differences in gene expression levels, mutational burden, copy number alterations (CNAs), and ligand-receptor interactions between short and long survivors, suggesting that strict treatment outcome-based stratification can enhance understanding of the variation between tumors and improve prediction of treatment response^10–12^.

At the molecular level, HGSC is characterized by genetic instability due to functional loss of TP53, and is accompanied by HRD in approximately 50% of patients^13,14^. HGSC also exhibits high levels of inter- and intra-patient heterogeneity, low frequency of recurrent oncogene mutations, and extensive CNAs^15,16^. The high levels of heterogeneity have made it difficult to identify reproducible prognostic markers^17,18^. To exploit the heterogeneity, a deeper understanding of its molecular features is required. While single-cell sequencing has increased knowledge of the tumor microenvironment and cellular composition of HGSC, it ultimately lacks the spatial context required to more completely comprehend the tissue biology^12,19^. Spatial transcriptomics has emerged as a transformative technology that enables the measurement of gene expression within the spatial context of the tissue architecture, allowing the investigation of cellular interactions, tumor microenvironment dynamics, and intratumoral heterogeneity^20–22^. Inference of genetic alterations, including CNAs, point mutations, and allelic imbalances directly from spatial transcriptomic data further provides new insights into tumor evolution and heterogeneity^23–26^. Spatial approaches have recently been applied to HGSC at cohort scale. A single-cell spatial atlas using independent HGSC tumors and targeting a preselected antibody panel at single-cell resolution showed that immune cell coinfiltration at the tumor-stroma interface associates with overall survival^27^. Transcriptome-wide spatial profiling of matched treatment-naive adnexal and omental tumors, stratified by progression-free interval, complements such large-scale atlas efforts.

In this study, we performed Visium spatial transcriptomic profiling of paired treatment-naïve adnexal and omental HGSC samples. We asked whether outcome-associated transcriptional programs were reproducible across anatomic sites, and whether they localized specifically to the tumor epithelium or stroma. Pathway and differential-expression analyses, integrated with copy-number inference, identified several candidate genes associated with unfavorable outcome. We validated their expression at the tissue level using RNA in-situ hybridization, and evaluated their prognostic associations in external cohorts. Finally, we explored whether the outcome-associated phenotype could occur as a spatially distinct, copy-number-defined subclonal state within an individual tumor.

## Results

### Matched adnexal and omental spatial profiling across treatment outcome groups

To compare adnexal and omental sites within the same patient, we assembled a cohort of nine individuals with treatment-naïve, advanced HGSC (FIGO stages III–IV) from whom both an adnexal and a matched omental tumor sample were available (Figure 1). All patients underwent primary cytoreductive surgery followed by platinum-based chemotherapy, and were stratified by platinum-free interval (PFI) into short (< 6 months), medium (6–18 months) and long (> 18 months) outcome groups (Table 1). Because both tumor sites are derived from the same patient at the same time, site-associated and outcome-associated signals can be separated within a single cohort.

**Table 1.** Summary of clinical data.

| Sample ID | FIGO stage | R status (mm) | Sample collection | Chemotherapy cycles | Survival | PFI days | PFI group |
| --- | --- | --- | --- | --- | --- | --- | --- |
| H110 | IIIC | 0 | PCS | 6 | Alive | 2134 | Long |
| H213 | IIIC | 0 | PCS | 6 | Alive | 1611 | Long |
| H119 | IVB | > 10 | PCS | 9 | Dead | 568 | Long |
| H180 | IIIC | 1-10 | PCS | 6 | Dead | 341 | Medium |
| H197 | IVA | 1-10 | PCS | 6 | Dead | 280 | Medium |
| H233 | IVA | 0 | PCS | 6 | Dead | 184 | Short |
| H178 | IVB | > 10 | PCS | 6 | Dead | 174 | Short |
| H152 | IIIC | 1-10 | PCS | 6 | Dead | 154 | Short |
| H145 | IIIC | 1-10 | PCS | 6 | Dead | 116 | Short |

FIGO, International Federation of Gynecology and Obstetrics; PCS, primary cytoreductive surgery; PFI, platinum-free interval; R status, residual tumor

**Figure 1.**
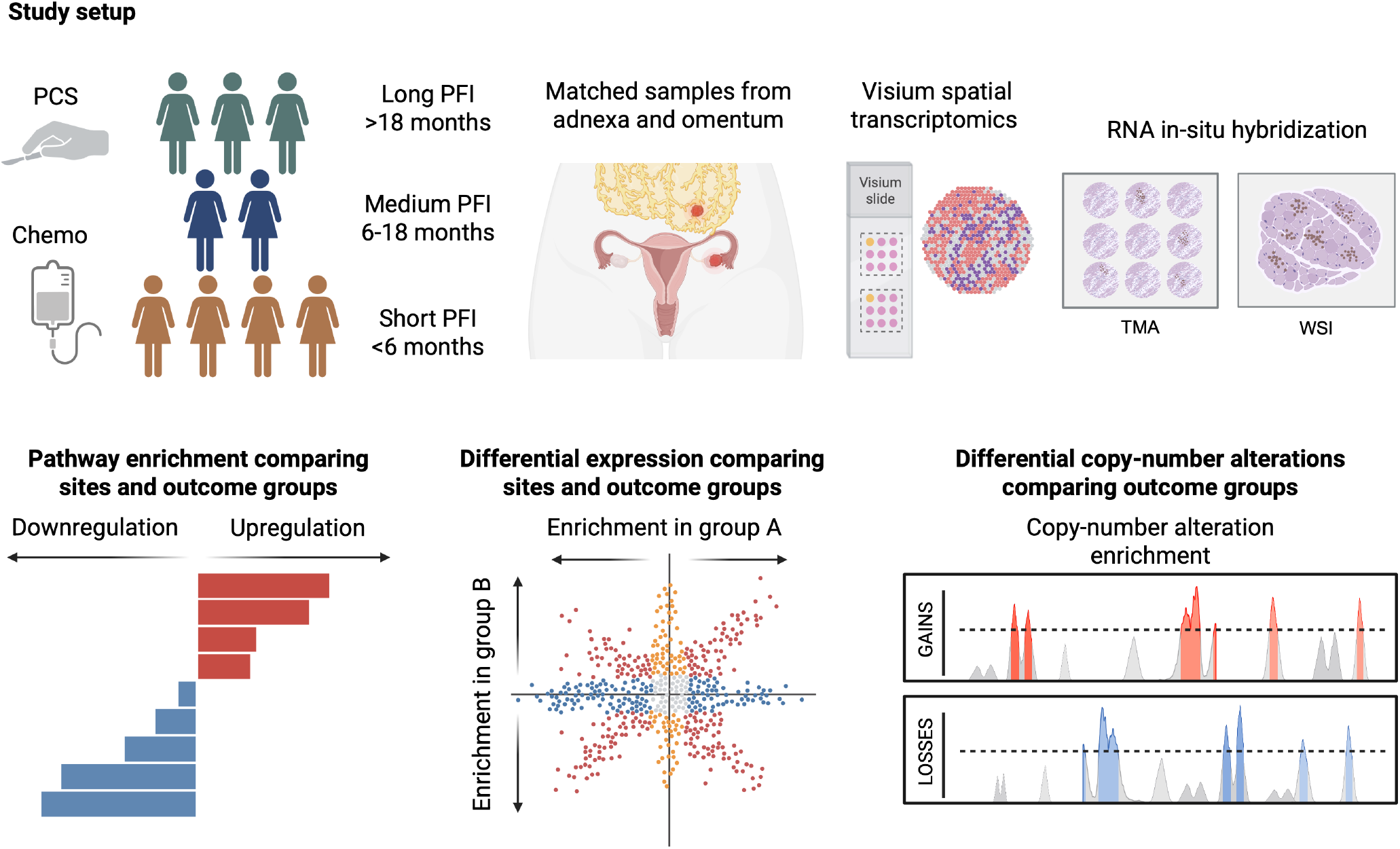
Patient cohort and study design. Top left: patient cohort description (n=9). Top middle: tumor sample collection sites. Top right: tumor profiling and validation methods. Bottom: computational methods used. PCS, primary cytoreductive surgery; Chemo, chemotherapy; PFI, platinum-free interval; TMA, tissue microarray; WSI, whole-slide image.

Spatial transcriptomic profiling using Visium for FFPE v2 of the 18 samples yielded, after quality filtering, 7,114 spots, of which 2,910 were annotated by a pathologist as tumor epithelium, 2,005 as stroma (and other) and 2,199 as mixed. The UMAP projection separated spots by individual, and within individuals by tumor site, consistent with the pronounced inter- and intratumoral heterogeneity described in HGSC (Supplementary Figure 1). Cell types were resolved within each spot by deconvolution against an external ovarian cancer single-cell reference^28^; estimated cell type proportions did not differ detectably between PFI groups or tumor site (Supplementary Figure 2b-d).

### Pathway enrichment analysis in the context of patient outcome

To assess whether biologically meaningful differences exist between outcome groups and tumor sites, we performed pathway enrichment analysis (GSEA) against MSigDB Hallmark collection, comparing the short PFI group to the pooled medium and long (med/long) PFI group (Figure 2a). Whole tissue core comparison revealed that the interferon alpha and gamma response pathways were the two most enriched gene sets in the short PFI group (NES = 2.68; NES = 2.57), accompanied by four additional inflammation-associated gene sets; coagulation, inflammatory response IL-6/JAK/STAT3 signaling and complement. The med/long PFI group was instead enriched by proliferative and cell-cycle regulatory programs (E2F targets, G2M checkpoints and MYC targets V1/V2) together with metabolic programs (mTORC1 signaling, unfolded protein response and glycolysis). Thus, the transcriptional distinction between outcome groups is characterized by an inflammatory profile in the short PFI group versus a proliferative and metabolic profile in the med/long PFI group.

**Figure 2.**
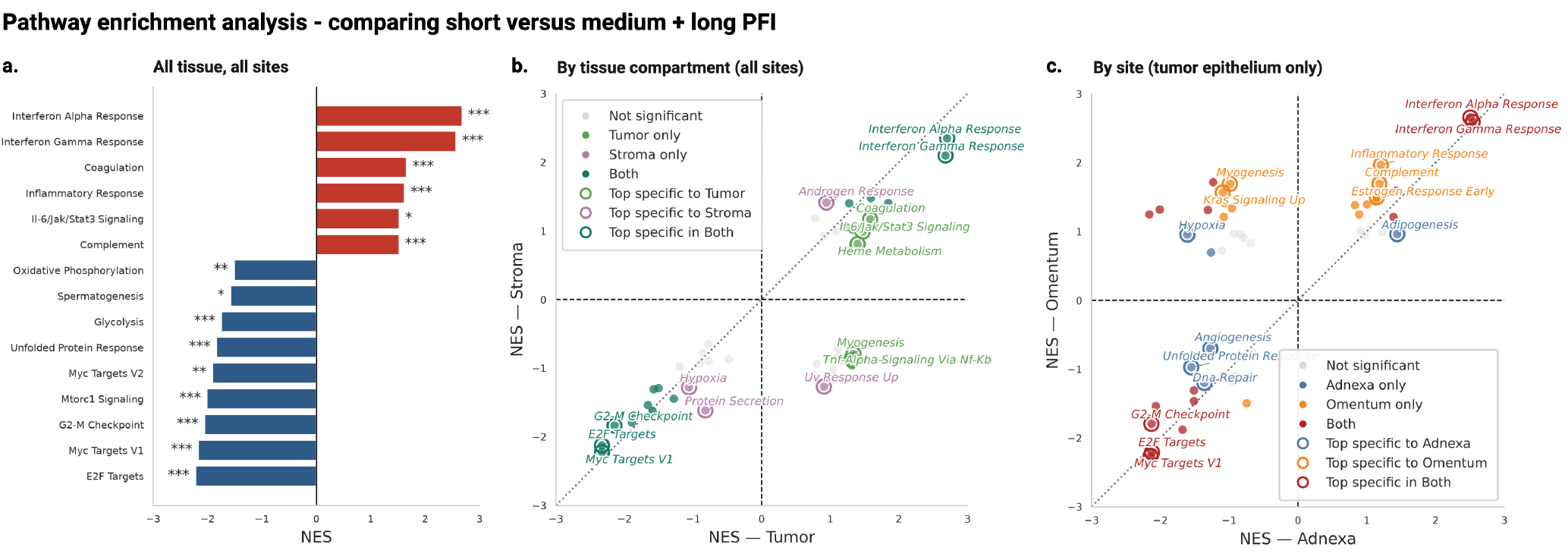
Pathway enrichment analysis by PFI group. **a.** GSEA against the MSigDB Hallmark (2020) collection across all samples, comparing the short PFI group with the pooled medium and long PFI groups. Positive NES (red) indicates enrichment in the short PFI group; negative NES (blue) indicates enrichment in the pooled medium and long PFI groups. *** p < 0.001; ** 0.001 < p < 0.01; * 0.01 < p < 0.05. **b, c**. Concordance of enriched pathways in the short versus pooled medium and long PFI groups between (**b**) tumor epithelium and stroma, and (**c**) adnexal and omental sites (within tumor epithelium only). Each point is one Hallmark gene set; axes show its NES for short versus pooled medium and long PFI within each of the two groups compared. Points are colored by FDR significance (q < 0.25) in each group: significant in the first group only (**b**, tumor epithelium, light green; **c**, adnexa, blue), in the second only (**b**, stroma, purple; **c**, omentum, orange), in both (**b**, dark green; **c**, red), or in neither (gray). The dotted diagonal marks concordant enrichment between the two groups; open circles mark the five most concordant and diverging gene sets from the diagonal respectively in each significance category, labeled by name. GSEA, gene set enrichment analysis; NES, normalized enrichment score; PFI, platinum-free interval; FDR, false discovery rate.

As hallmark inflammatory signatures can be driven by non-epithelial cells and may be further enriched at certain tumor sites, we asked whether the short PFI interferon effect was restricted by tissue compartment (Figure 2b) or tumor site (Figure 2c). Interferon alpha and gamma were significantly increased in both tissue compartments and at both tumor sites, indicating a tissue-wide inflammatory phenotype, that is neither solely an artifact of stromal admixture nor restricted to a single tumor site. Aside from consistent enrichment of interferon response, the other inflammation-related gene sets showed greater variation across tissue compartments and tumor sites in the short PFI group. The tissue compartment comparison revealed that IL-6/JAK/STAT3 signaling and coagulation were enriched in both compartments but reached significance only in tumor epithelium (Figure 2b, upper right quadrant). Similarly, complement and inflammatory response were enriched at both tumor sites but reached significance only in omentum (Figure 2c, upper right quadrant). Most of the remaining Hallmark sets fell near the diagonal; however, a number of diverging pathways showed opposing enrichment patterns across PFI groups. We note especially that the inflammation- and proliferation-linked TNF-alpha signaling via NF-kB was significantly enriched in tumor epithelium in the short PFI group and downregulated in stroma (Figure 2b, lower right quadrant). The short PFI signature at the omental site therefore carried a slightly broader inflammatory component as compared to the adnexal site.

In the med/long PFI group, MYC targets V1 was the most consistently upregulated gene set across tissue compartments and tumor sites, followed by cell-cycle regulatory gene sets E2F targets and G2M checkpoints (Figure 2a). The tumor site comparison further showed that while enriched in both, DNA repair, angiogenesis and unfolded protein response were significant only in adnexa (Figure 2c, lower right quadrant). In contrast, while hypoxia was enriched in both tumor and stroma in the med/long PFI group, the site comparison revealed significant enrichment specifically in adnexal tumor epithelium (Figure 2b, lower left quadrant) and downregulation in omentum (Figure 2c, upper left quadrant). These findings suggest that this proliferative and cell cycle regulatory program is a shared feature of the med/long PFI tumors, independent of tissue compartment or tumor site.

Since cancer-cell-intrinsic MHC class II expression has been identified as a leading predictor of prolonged survival in HGSC^27^, and MHC class II genes are members of the Hallmark interferon gamma response set, we asked whether the short PFI interferon enrichment reflected an antigen-presentation program or if it is carried more by effector interferon-stimulated genes like BST2 and IFI27 which we found to be enriched in the short PFI group in our cohort. Testing depletion of the antigen-presentation module against the remaining members of the IFN*γ* gene set present in our data gave a one-sided Fisher *p* = 0.021 (odds ratio 0.13) (see Supplementary Results). The short PFI interferon signature we observe is thus more like an effector interferon-stimulated gene program from which antigen presentation, and class II in particular, is mostly excluded.

Taken together, Figure 2 identifies the interferon response as the most consistent transcriptional correlate of short PFI in this cohort, reproducible across tissue compartments and across matched primary and metastatic sites, with complement signaling enriched alongside it. These pathway-level results directed the subsequent differential expression and copy-number analyses towards interferon-stimulated genes and complement components.

### Differential gene expression and copy-number analysis identify short-PFI associated gene candidates

Differential gene expression analysis was performed on patient level to detect specific gene candidates with distinct expression profiles in the outcome groups (short PFI versus med/long PFI), and in different tissue compartments (tumor epithelium, stroma, and mixed tumor epithelium+stroma) (Supplementary Figure 4). We then performed a marker gene analysis on the Visium spots annotated as tumor epithelium only, identifying multiple genes associated with each PFI group across tumor sites (Figure 3a). To identify the most clinically useful PFI-associated candidate biomarkers, we focused on DEGs which correlated with outcome in both adnexal and omental sites. Amongst the top candidates, we identified two genes, C3 and IFI27, with higher expression in short PFI samples at both tumor sites. MIF showed higher expression across tumor sites collectively in both medium and long PFI groups.

**Figure 3.**
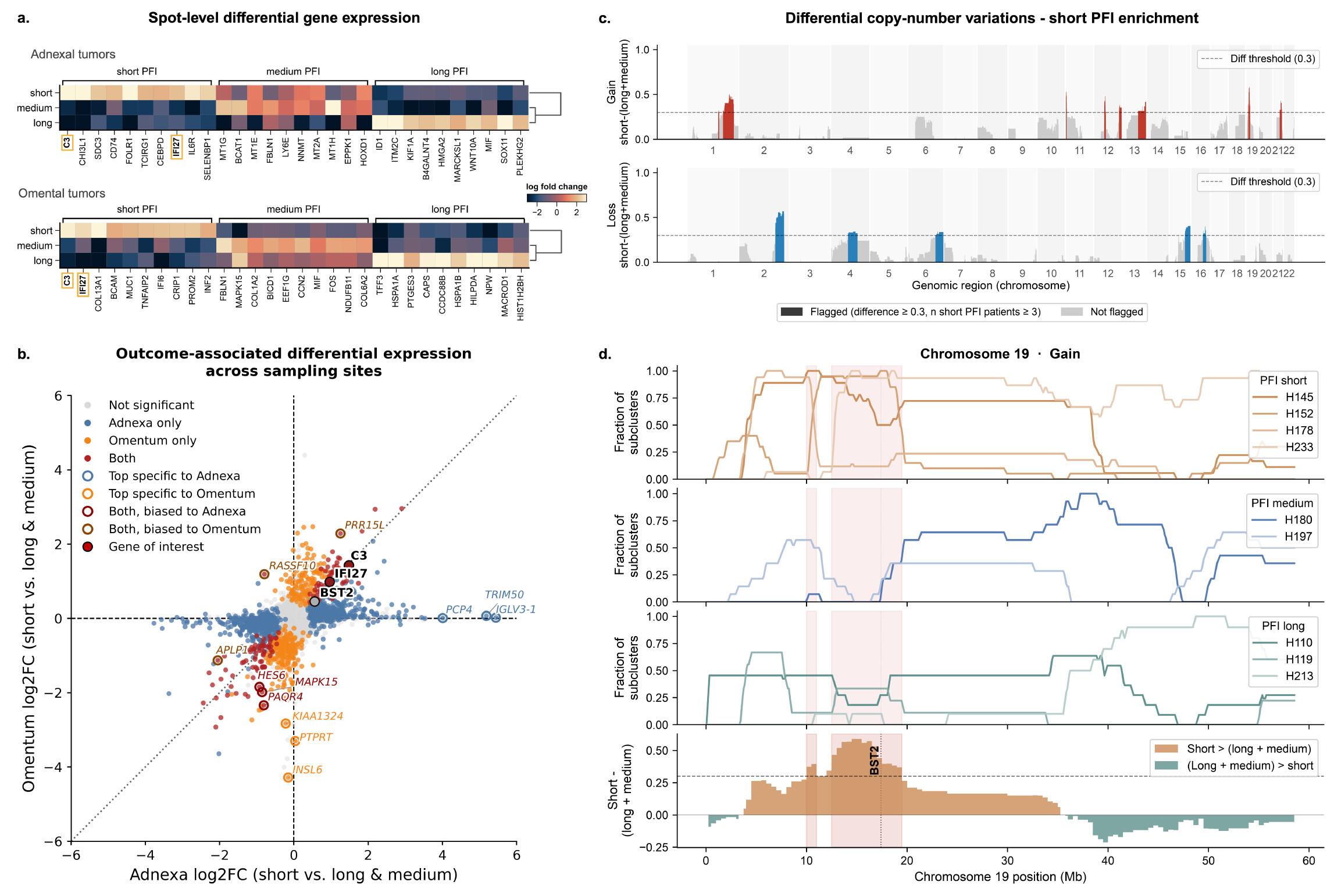
Differential gene expression and copy-number analysis identify short PFI associated gene candidates. a. Heatmaps illustrating the top ten differentially expressed genes by PFI group in tumor epithelium from adnexal tumors (upper) and omental tumors (lower). Genes common to both tumor sites in the short PFI group are boxed in orange and bolded. **b**. Differential expression analysis, contrasting genes associated with short PFI in adnexal and omental tumor sites, incorporating all the spots and tissue compartments. Genes selected for tissue-level validation are highlighted. **c**. Genome-wide differential copy-number burden in short PFI. Each bar represents a 1 Mb genomic bin. Bar height reflects the differential copy-number burden score, defined as the difference in mean subcluster gain fraction (top) or loss fraction (bottom) between short PFI and long+medium PFI patients. Bins enriched in short PFI patients are colored red (copy-number gains) or blue (copy-number loss); grey bins did not meet the enrichment threshold. The dashed horizontal line denotes the differential score threshold. **d**. Differential copy-number gains on Chromosome 19. Lines represent individual patients; orange fill indicates regions where short PFI exceeds long PFI, green fill the reverse. PFI, platinum-free interval; CNV, copy-number variation. Figure 6b&c). We also found a strong enrichment of BST2 in the tumor epithelial subclone s1, which harbors a similar Chromosome 19 amplification (Figure 6c).

To identify genes associated with short PFI across tumor sites, we performed a cross-site differential expression analysis, accounting for inter-patient variability, and incorporating all tissue compartment annotations. These findings confirm the previously flagged gene candidates C3 and IFI27 as significantly associated with short PFI in both adnexal and omental tumor sites. Marker genes specific to adnexal samples included IGLV3-1, TRIM50, and PCP4, upregulated in the short PFI group, as well as genes INSL6, PTPRT, and KIAA1324, which were specific to omental samples and downregulated in short the PFI group. (Figure 3b).

In parallel, given that HGSC is a copy number-driven cancer, we performed a genome-wide copy-number analysis to independently assess whether genomic alterations differed between PFI groups. Copy-number variations (CNVs) were inferred from Visium spots annotated as tumor epithelium and mixed tumor epithelium and stroma using InferCNV. Spots annotated as stroma, from all samples collectively, were used as reference. The overall CNV profile revealed significant inter-patient heterogeneity, with intra-patient heterogeneity observed to a lesser extent (Supplementary Figure 3). No significant association between PFI and the number, length or percentage genome altered *derived from the CNVs*, was observed. We therefore shifted our focus to regions of the genome that showed alterations specifically enriched in the short PFI group. To identify these regions, we developed a differential CNV score (diff-cnv score) to quantify the enrichment of CNVs in short versus medium and long PFI groups (see methods). The full differential CNV profile can be seen in (Figure 3c).

The largest differential CNV loss (0.6) was observed on chromosome 2, spanning a region of 43 Megabases (Mb), containing both STAT1 and STAT4, key mediators of interferon signaling (Supplementary Figure 5). As STAT1 is a known driver of MHC class I and II antigen presentation^29^, its loss supports the impaired antigen presentation observed in short-PFI tumors. Among copy-number gains, the longest differential CNV was observed on chromosome 1, with a gain over a stretch of 53 (diff-cnv score of 0.5; Supplementary Figure 6). Chromosome 19 harbored the most prominent differential copy number gain (diff-cnv score of 0.589), affecting a genomic stretch of roughly 7 Mb in size from base 10, 250, 000 through 19, 500, 000. Notably, the chromosome 19 gain region contained BST2 which is an interferon-stimulated gene (Figure 3d). Together, based on differential gene expression, CNVs, and pathway enrichment findings, we selected C3, IFI27 and BST2 for tissue-level validation.

### Tissue level validation of genes correlated with short PFI in spatial transcriptomics

To validate the Visium ST findings, we performed RNA in-situ hybridization (RNA-ISH) for C3 (complement component 3), as well as interferon response pathway associated genes IFI27 (interferon alpha induced protein 27) and BST2 (bone marrow stromal antigen 2). C3 and IFI27 were selected for validation based on their significant upregulation in the short PFI group as compared to medium and long PFI. BST2 was chosen specifically due to copy-number gains detected on chromosome 19 in short PFI patients (Figure 3). In ST, C3 was highly expressed in short PFI tumor epithelium but also in mixed regions when compared to medium and long PFI groups. RNA-ISH tissue level validation, performed on subsequent sections from the same blocks used for Visium spatial transcriptomics, matched these ST results. C3 expression was significantly higher in short PFI tumor epithelial compartment, with lower scores in medium and long PFI groups (Figure 4a).

**Figure 4.**
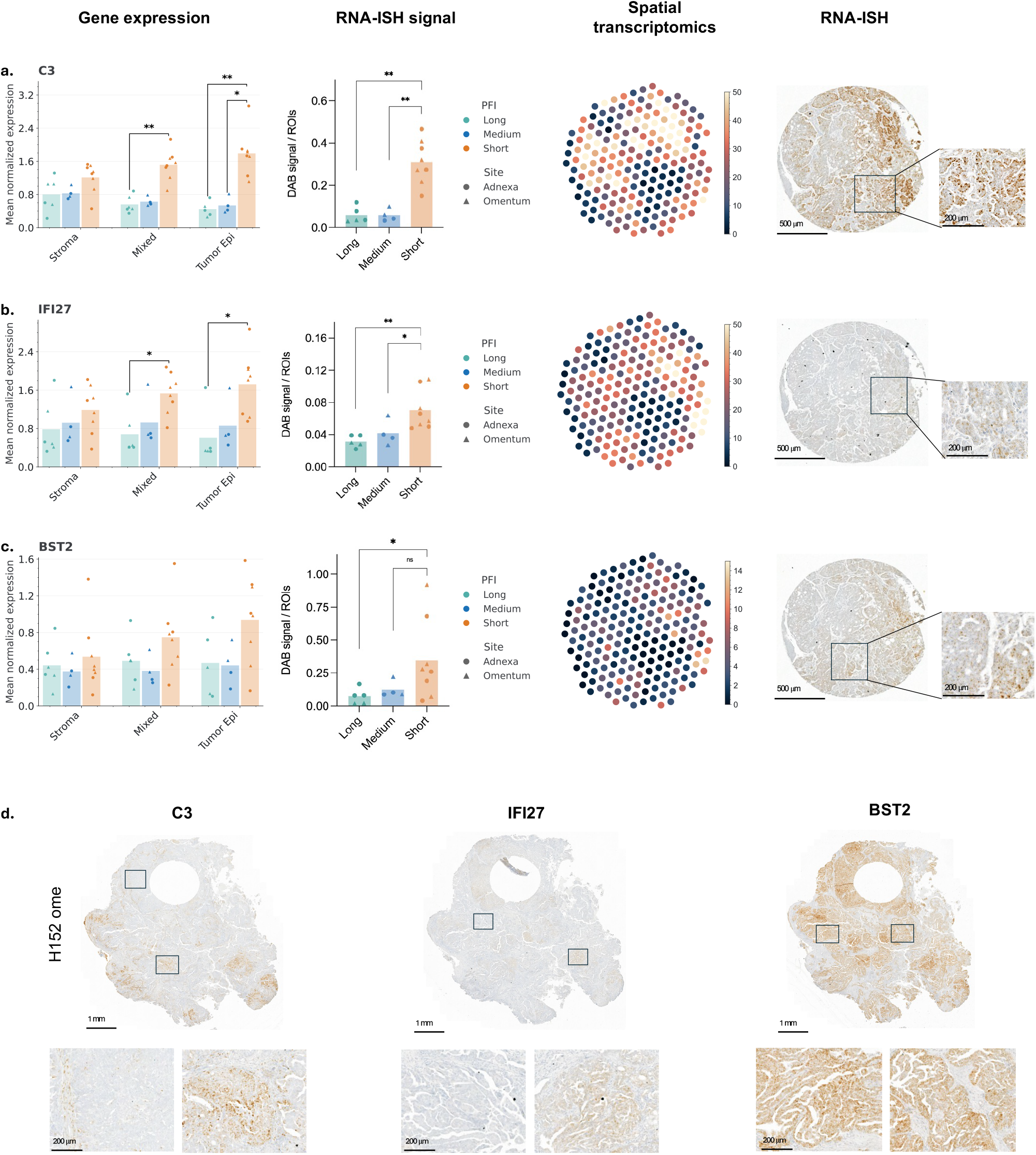
Spatial expression and validation of selected genes. a-c. From left to right: normalized ST gene expression by tissue type and patient outcome, RNA-ISH scores from tumor epithelium by patient outcome, ST visualization, and RNA-ISH. for C3 (**a**.), IFI27 (**b**.) and BST2 (**c**.). In the first two columns, colors indicate PFI groups and shapes represent the mean expression per patient and sampling site. In columns three and four, the spatial visualizations are derived from patient H178. **d**. Representative whole slide images from one patient (H152, omentum) showing the expression pattern of C3, IFI27 and BST2 as detected by RNA-ISH. ST, spatial transcriptomics; RNA-ISH, RNA in-situ hybridization; PFI, platinum-free interval. Significance is indicated as asterisks: *** *p <* 0.001; ** *p <* 0.01; * *p <* 0.05; ns, not significant

IFI27 followed similar pattern in ST and was found to be highly expressed in short PFI tumors, especially in tumor epithelium and mixed compartments; RNA-ISH validation further confirmed significantly increased expression in tumor epithelium. BST2, by contrast, was more specific to tumor epithelium in ST, though its upregulation did not reach significance at the patient level. Tissue-level validation, however, successfully distinguished short from long PFI group (Figure 4b-c).

Furthermore, to assess whether TMA-based findings can be recapitulated across the tumor in a broader region, we performed RNA-ISH on five whole tissue sections, including two short PFI patients (matching adnexal and omental tumors) and one medium PFI patient (omental tumor). C3 and IFI27 showed heterogeneous expression patterns with variable intensity in both tumor and stromal regions in short PFI patients. BST2 was diffuse throughout the tumor and variable in stromal regions in short PFI. All three genes showed low/no expression in the medium PFI patient sample, in line with the TMA findings (Figure 4d).

### Survival analysis in external cohorts

To further validate our findings on C3, IFI27 and BST2 we evaluated their prognostic potential using external publicly available ovarian cancer cohorts, TCGA-OVCA (n = 489)^30^ and GSE9891 (n = 285)^31^. Of the three, only C3 was consistently associated with clinical outcome. We found that high C3 expression was significantly associated with short progression-free survival (PFS) in both external ovarian cancer cohorts (p = 0.048 and p = 0.0083, respectively). These findings align with both our ST data and tissue level validation where C3 showed the strongest correlation between high expression and short PFI. While the TCGA cohort showed weak but significant (p = 0.039) link between high C3 expression and overall survival it could not be replicated in GSE9891 cohort (Figure 5). Neither IFI27 or BST2 was found to correlate with PFS or overall survival in the external cohorts. Taken together, these findings identify C3 as the most clinically interesting candidate, showing consistent association to short PFI across discovery and external validation cohorts, while the prognostic value of IFI27 and BST2 may be better captured by ST rather than bulk expression data.

**Figure 5.**
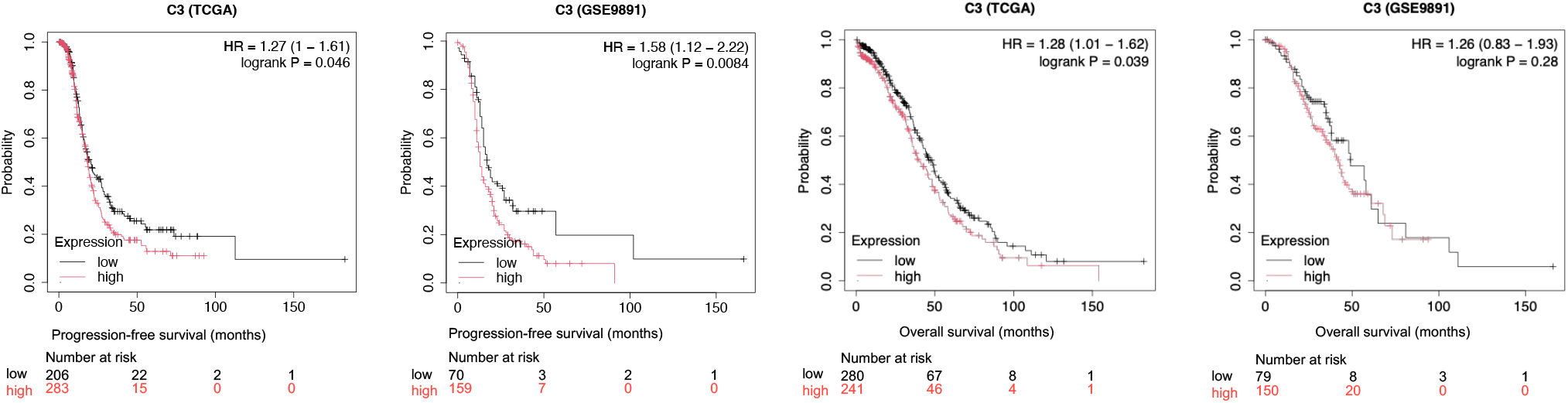
Correlation of C3 with survival. Progression-free survival and overall survival were compared between high and low C3 expression. Kaplan-Meiers were created using KMplotter (https://kmplot.com/analysis/).

### Subclonal heterogeneity of the interferon/complement phenotype: a case study in an independent cohort

Our primary cohort analyses treated PFI-associated gene expression as a patient- or tumor-site-level property. However, HGSC is known to exhibit substantial intratumoral heterogeneity, raising the question of whether the interferon/complement phenotype identified above is uniformly present within a tumor or itself varies at the subclonal level. To explore this, we examined copy-number-defined subclones within a single short PFI adnexal tumor (OVA09) from an independent, previously published HGSC cohort^32^. This cohort was collected after primary debulking surgery, as in the present study, but contains only adnexal samples; no omental comparison is possible in this analysis. CNVs were inferred using the same approach as applied to the primary cohort, revealing a similar degree of inter-patient heterogeneity (Supplementary Figure 7).

Within a single TMA spot from OVA09, we identified four spatially distinct subclones in areas annotated as tumor stroma, each characterized by a unique CNV profile. The patient-matched tumor epithelium showed three relatively uniform CNV subclones (Figure 6a-b). Shown in Figure 6a, middle panel, are only areas of the TMA that contained tumor epithelium and tumor stroma as per pathologist annotations. All other TMA spots from this cohort showed homogeneous CNV profiles. Genes identified to be associated with short PFI in the primary cohort were found upregulated in the subclones of OVA09 (Figure 6c). Notably, BST2, C3 (Chromosome 19), and IFI27 (Chromosome 14) were enriched in the subclone (s4) that also harbors a Chromosome 19 copy-number gain similar to that identified in short the PFI group in our primary cohort (Figure 3,

**Figure 6.**
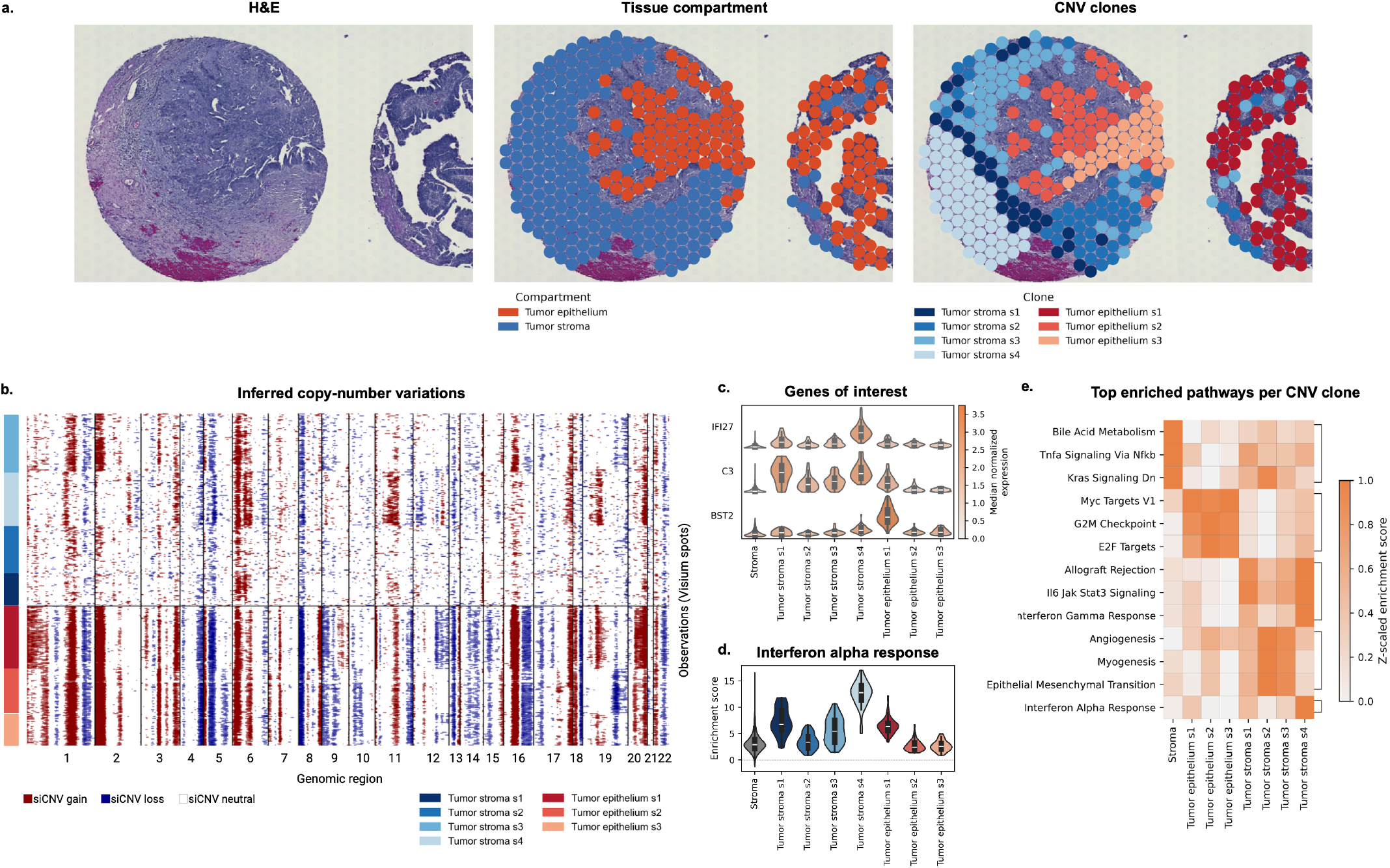
Copy-number clonality in the tumor-stroma border. a. Tissue morphology, annotated compartments, and CNV clones in short PFI patient OVA09. **b**. CNVs were inferred using InferCNV from areas annotated as tumor stroma and tumor epithelium by a pathologist. Ovarian stroma from all patient’s TMAs was used as a reference for InferCNV. **c**. Normalized median expression of IFI27, C3, and BST2. **d**. Highlighted interferon alpha response pathway enrichment score across all CNV clones. **e**. Top enriched pathways from the cancer hallmarks collection per CNV clone. CNV, copy-number variation

Gene set enrichment analysis further showed that the IFN-*α* and -*γ* response pathways were significantly upregulated in subclone s4 relative to subclones s1-s3 and stroma (Figure 6d-e). This pattern is compatible with the enrichment observed in the short-versus-long PFI comparison in our primary cohort, raising the possibility that the interferon-high phenotype associated with short PFI at the patient level can also emerge as a distinct subclonal population within a single tumor. By contrast, subclones s2 and s3 showed upregulation of the angiogenesis, myogenesis, and epithelial mesenchymal transition pathways, pointing to a distinct transcriptional program coexisting within the same tumor.

As this analysis is based on a single patient and a single annotated region, it should be interpreted as a hypothesis-generating case study rather than a replication of the primary cohort findings.

## Discussion

Currently available clinical tools cannot reliably identify which patients with HGSC are most likely to experience earlier disease relapse, highlighting the need for new approaches. In this study, we used spatial transcriptomics to investigate whether features of the tumor and its microenvironment are associated with treatment outcome, taking advantage of an advantageous design incorporating patient-matched adnexal and omental tumors. Our findings identify a distinct molecular phenotype associated with more rapid disease recurrence (short PFI), characterized in part by complement and interferon-related signaling. Among the candidate genes identified, C3 showed the strongest clinical relevance, with its association with poor outcome further supported in external cohorts. We also uncovered unexpected spatial genomic heterogeneity within the tumor microenvironment, including copy-number alterations in regions classified as stroma. Together, these findings highlight spatially resolved molecular features of HGSC that may improve our understanding of treatment response and provide new directions for predictive biomarker development.

Tumor-intrinsic inflammatory signaling is increasingly recognized as a contributor to disease progression and treatment resistance in HGSC. Consistent with a role for immune signaling in HGSC outcome, intratumoral CD8^+^ and CD4^+^ T cell infiltration is a well-established predictor of improved survival, an axis sustained in part by IFN-*γ*-dependent chemokine production and antigen presentation^11,33^. A recent single-cell HGSC study has shown that complement and interferon signatures are highly correlated within the same tumor cell populations, and are already present in treatment-naïve samples, suggesting a shared, chronic inflammatory state rather than two distinct processes^34^. Our findings build on this as we show that enhanced complement and interferon signaling are features of all tissue compartments and tumor sites in the short PFI group. Medium and long PFI tumors instead showed enrichment in metabolic and proliferative programs rather than inflammatory ones. A negative correlation between inflammatory vs metabolic/proliferative programs has been previously suggested^34,35^. Our ST analysis also showed consistently higher expression of C3, a key complement gene, in tumor, stroma and mixed regions. Whole-slide validation revealed a spatially heterogeneous staining pattern, emphasizing region-specific rather than tumor wide expression. In HGSC, C3 has been reported to promote ovarian cancer proliferation and to drive tumor progression especially in metastatic tumors^36,37^. A recent spatial transcriptomics study (n = 8) showed that high tumor-derived C3 was associated with recurrent HGSC, further strengthening the link to poor outcome^38^. Whole-slide validation of interferon-stimulated genes further revealed that IFI27 expression was heterogeneous across both tumor and stromal regions while BST2 remained more specifically tumor derived. Notably, BST2 also showed copy number gains in short PFI samples, suggesting that its upregulation might be partly driven by gene amplification. These patterns were consistent with our ST findings and suggest that while both genes are interferon-driven, IFI27 may reflect a more tumor and tumor microenvironment wide signal whereas BST2 indicates a more tumor-intrinsic response. In ovarian cancer, IFI27 has been linked to cisplatin resistance and poor disease-free survival, while BST2 has been shown to promote an immunosuppressive tumor microenvironment, driving disease progression^39–41^. Together, these findings suggest that pre-existing complement/interferon phenotype contributes to poor treatment response in HGSC.

The association of higher interferon signaling and shorter PFI observed in our cohort requires careful interpretation, as interferon signaling can have opposing effects in chemotherapy outcome in HGSC. A previous multi-omics study on chemo-refractory HGSC showed that reduced baseline type I interferon activity is associated with primary chemoresistance.^42^. Consistent with this, cancer-cell-autonomous type I interferon signaling has been found to enhance the efficacy of chemotherapy as well as predict the treatment response in breast cancer cohorts^43^. Our observation is not directly comparable as those analyses stratify on primary refractory disease, whereas our individuals all completed platinum-based therapy and are stratified by the platinium-free interval. Additionally, our enrichment is measured as Hallmark interferon-response gene-set activity within spatially resolved epithelial regions rather than as pathway activity across a whole biopsy. Conversely, persistent or chronic interferon activation has been linked to immune evasion and shown to drive platinum resistance in HGSC cell models^44,45^. Interferon signaling can also induce MHC-II expression which has been linked to better prognosis and higher immune cell infiltration in several cancers^29,46^. A recent paper on HGSC likewise reported cancer-cell MHC-II expression as a favorable marker of overall survival^27^. Our data are consistent with this direction rather than in conflict with it, as the interferon program enriched in short PFI tumors is composed of effector interferon-stimulated genes (ISGs), including IFI27 and BST2, and specifically excludes antigen presentation. As sustained tumor-cell derived ISGs expression is associated with treatment resistance, the repression of MCH-II may further limit tumor cells recognition from immune system enhancing immune evasion and tumor progression. Hence, we suggest that interferon activation and antigen presentation should be treated as separate axes. Further studies are needed to explore how pre-existing versus acute interferon signaling relates to antigen presentation within the same spatial compartments.

To further validate our findings, we evaluated the prognostic potential of C3, IFI27 and BST2 using publicly available ovarian cancer cohorts. Only high C3 expression was significantly associated with shorter progression-free survival, aligning with our spatial and tissue-level findings linking high C3 expression to short PFI. The lack of replication for IFI27 and BST2 may partly reflect the composition of bulk RNA-seq data which averages signal across mixed tissue and cell types rather than capturing tissue type expression. C3, by contrast, showed a robust association with outcome across spatial, tissue-level, and bulk analyses, highlighting its relevance as a candidate therapeutic target. Notably, Pegcetacoplan, a C3 inhibitor, is currently being tested in a Phase 2 trial in recurrent ovarian cancer patients (NCT04919629) with preliminary results showing an early signal of clinical benefit^47^.

Our case-study analysis of copy-number-defined subclones in an independent HGSC cohort suggests that this interferon/complement phenotype is not necessarily a uniform property of a tumor, but can instead emerge as a distinct subclonal population coexisting alongside subclones with other transcriptional programs, such as angiogenesis. This observation, while based on a single patient, is consistent with the broader recognition that HGSC subclones can differ substantially in their immune and inflammatory profiles, and raises the possibility that the interferon/complement-high phenotype we associate with short PFI arises focally within a tumor before becoming dominant, rather than developing uniformly across all tumor cells. If confirmed in larger cohorts, these findings would suggest that broader spatial sampling may be required to reliably detect focal interferon/complement-high subclones associated with short PFI.

Several limitations should be considered when interpreting these findings. In particular, our cohort is small, which limits statistical power and precludes multivariable adjustment; group comparisons are also sensitive to individual patients. To mitigate this, we derived pathway enrichment and DEGs from patient-level pseudobulk profiles rather than from individual spots, so that patients, rather than spots, constitute the unit of analysis. To maximize the number of patients and samples profiled, we utilized TMAs rather than single whole tissue sections. Consequently, only two regions of tumor were sampled from each patient, which cannot capture the full extent of intratumoral heterogeneity. To address this, we evaluated the data in different tissue compartments, and included whole tissue sections in the RNA-ISH validation to visualize the potential heterogeneity of our markers. In reality, even a whole tissue section is very likely to underrepresent heterogeneity in this tumor type, given the range of tumor sites and sampling styles. Nonetheless, these outcome-stratified data provide valuable hypothesis-generating associations which warrant validation in larger series.

In conclusion, we have shown that interferon signaling is associated with short PFI across matched adnexal and omental HGSC tumors, identified C3 as a candidate prognostic biomarker, and found preliminary evidence of copy-number variation in HGSC tumor stroma. Future studies should expand the number of spatially profiled HGSC samples, as spatial analyses can provide additional insight into the molecular biology of HGSC and inform clinical decision-making for the benefit of patients. As the number of spatial transcriptomic technologies continues to increase, additional knowledge from whole-transcriptome, single-cell spatial approaches should be integrated. Finally, efforts should focus on moving from the identification of clinically relevant signals toward their validation, translation and clinical implementation.

## Methods

### Ethical permission and clinical data

The study protocol is approved by the Hospital District of Southwestern Finland Ethics committee (ETMK Dnro: 145/1801/2015) and is part of HERCULES/DECIDER ovarian cancer research project (ClinicalTrials.gov identifier NCT04846933). All study participants have given their written consent prior to enrollment. High-grade serous ovarian carcinoma tumor samples from 9 individuals with stage III-IV disease at diagnosis were included. Detailed inclusion criteria is shown in Supplementary figure 8. All individuals underwent primary cytoreductive surgery, followed by 6-9 cycles of platinum-based chemotherapy. Individuals were categorized based on the platinum-free interval (PFI; short < 6 months, medium 6-18 months, and long > 18 months). PFI is defined as the length of time between the last dose of platinum and the date of recurrence. Both adnexal and omental samples were collected from each study participant during primary cytoreductive surgery.

### TMA construction

Tissue was preserved using PAXgene fixation, following the manufacturer’s protocol (#765312, Qiagen), and stored at 4 °C. The tumor content of each block was assessed with haemoxylin and eosin (H&E) by a pathologist. To assess tissue quality, two 10 um rolls were cut from each block and RNA isolated using QIAsymphony RNA kit (#931636, Qiagen). The DV200 value was measured using the Agilent TapeStation 4200 at the Functional Genomics Units at University of Helsinki and only samples with DV200 > 60% were included.

Tissue microarrays (TMAs) were created using H&E stained whole-slide images to identify the most representative tumor regions. One to two 1.5 mm tumor cores were obtained from each tissue block, based on the amount of tumor present, with a goal of four per patient (two adnexal, two omental). The final number of cores included per individual were as follows: six individuals had two adnexal cores and two omental cores, two individuals had two adnexal cores and one omental core, and one individual had one adnexal core and two omental cores . In total, four TMAs (6,5 mm x 6,5 mm) were constructed, each containing eight tumor cores and one orientation core (low-grade serous carcinoma) in 3×3 configuration. The TMAs were constructed using TMA Grand Master (3DHistech) and manual core boring at Helsinki Biobank.

### Visium spatial transcriptomic profiling

Spatial gene expression was studied using the Visium v2 Spatial Gene Expression kit for FFPE samples from 10X Genomics (#PN-1000188). In short, one 5 *µ*m section from each TMA was placed per capture area, followed by H&E staining. The slides were further processed according to the CG000407 Visium Spatial Gene Expression user guide for FFPE at FIMM Single-Cell Analytics unit at University of Helsinki. Optimal conditions were assessed before using the Tissue Section Test Slide kit (#PN-1000347) on a separate high-grade serous ovarian carcinoma tissue sample and the permeabilization time was set to 60 minutes. Libraries were sequenced to the depth of 100,000 reads per spot on the Illumina NovaSeq 6000 system using the recommended 28-10-10-50 read setup. Hematoxylin and eosin stainings of the tissue sections were performed and evaluated by a pathologist to confirm tissue type and tumor content.

### Visium data pre-processing

The raw data were processed with the SpaceRanger pipline (*v*2.0.0). FASTQ files were generated using the pipeline’s *bcl2fastq* wrapper function *spaceranger mkfastq*. From these FASTQ files, gene-barcode count matrices were generated using the GRCh38-2020-A as a reference. Resulting count matrices from the four capture areas were sequencing-depth normalized and combined into a single matrix using *spaceranger aggregate*.

Visium spatial transcriptomics data were processed as AnnData objects using SquidPy^48^ (*v*1.8.3) and Scanpy^49^ (*v*1.12.3) in Python. Spatial spots were assigned to outcome groups based on the associated clinical metadata. The data were filtered to include only spots with a minimum of 250 unique molecular identifiers (UMIs) and *>* 50 unique genes. Genes were filtered to retain only those expressed in at least three spots. Filtered data were subsequently count-normalized to a target sum of 10, 000 before applying the natural logarithm.

The H&E images were annotated by a trained pathologist (A.L.) using Loupe Browser (*v*.8.1.2, 10X Genomics). The tissue regions were categorized as “tumor”, “mixed” or “stroma”. Only spots fully covered by either tumor epithelium or stroma were annotated as such regions. Spots containing both tumor and stroma were annotated as mixed.

### Differential gene expression analysis

Raw counts from filtered data were aggregated into pseudobulk counts binning by platinum-free interval (PFI), tumor compartment as annotated by the pathologist, sampling site, and patient. Differentially expressed genes (DEGs) were identified using PyDESeq2^50^ (*v*0.5.4) on tissue compartment-, sampling site-, and outcome group-level, always comparing short PFI to long and medium combined. For visiualizations, log2 foldchange (log2fc) shrinkage was performed to stabalize results. To obtain spot-level DEGs for comparisons within the tumor epithelial cells only for Figure 3, log-transformed data were input into the scanpy function *rank_genes_groups*. DEG expression test was tested for using a Wilcoxon test. Resulting p-values were corrected for multiple-testing using Benjamin-Hochberg. Genes were considered differentially expressed when their adjusted p-value was *<* 0.05. Gene expression differences between PFI groups and tissue compartments for Figure 4 were determined by a Kruskal-Wallis test on patient-level pseudobulked data, followed by a Dunn’s post-hoc test for the subset analysis comparing the different tissue compartments as subgroups.

### Pathway analysis

Gene set enrichment analysis (GSEA) was performed using the Python package decoupler^51^ (v1.7.0). For all analyses, the Hallmark gene set collection from the Molecular Signatures Database^52^ (MSigDB) 2020 was used. The collection was filtered to include only gene sets with more than 15 and less than 500 genes. As input to decoupler, a list of genes ranked by differential gene expression using PyDESeq2 for short PFI based on the DEG analysis described above was used. The analyses are based on 1,000 random permutations, and the resulting enrichment scores were normalized, accounting for the gene set size and multiple testing.

To determine whether the interferon enrichment was carried by antigen presentation or by effector interferon-stimulated genes, antigen-presentation genes were defined as the classical and non-classical HLA class I and class II loci together with B2M, the peptide-loading machinery (TAP1, TAP2, TAPBP), the immunoproteasome subunits PSMB8 and PSMB9, and the class I and class II transactivators NLRC5 and CIITA, and subset to genes present in the measured Visium v2 probeset. Leading-edge membership of this module was compared with that of the remaining set members by one-sided Fisher exact test.

### CNV inference

Copy-number variations were inferred from the gene expression data using InferCNV^53^. As reference sets for all analyses, combined benign stroma from all patients from the corresponding cohort was used. Filtered count data were input to InferCNV. All inferCNV runs were performed using the following parameters unless otherwise specified: *cutoff=0*.*1, denoise=TRUE, HMM=TRUE*, and otherwise default parameters.

InferCNV was run in i6 HMM mode, which outputs predictions on six copy-number states (state 1: complete loss; state 2: heterozygous loss; state 3: neutral; state 4: gain; state 5: amplification; state 6: high-level amplification). InferCNV assigns HMM states to genomic regions within transcriptionally defined subclusters of spots; all downstream analyses were performed at the subcluster level and subsequently aggregated to the patient level as described below.

#### Genome-wide differential CNV score

To quantify copy-number burden across the genome in a patient-level, PFI-group-aware manner, a sliding-window differential CNV score (diff-cnv score) was developed. The genome was tiled into non-overlapping 1 Mb bins. For each bin, each chromosome, and each CNV type (gain: state*≥* 4; loss: state *≤*2) independently, we computed a per-sample gain (or loss) fraction, defined as the proportion of a patient’s inferCNV subclusters carrying the relevant CNV state in that bin. For gains, the maximum state across overlapping region entries was retained; for losses, the minimum state was retained. This yielded, for each chromosome and CNV direction, a bin × subcluster state matrix. For each sample *s* and bin*× b*, the alteration fraction was defined as the fraction of that sample’s subclusters carrying the alteration at that bin. For gains,

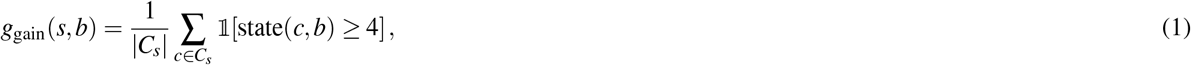

and for losses,

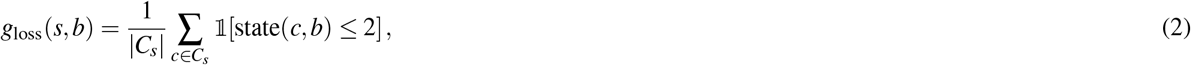

where *C*_*s*_ is the set of subclusters belonging to sample *s* and 1[*·*] is the indicator function. Both fractions lie in [0, 1], with 0 meaning no subcluster and 1 meaning all subclusters of the sample carry the alteration at bin *b*. In the following, *g*(*s, b*) denotes the relevant fraction for the CNV direction being scanned.

Samples were stratified by progression-free interval (PFI) category (short, medium, long). The *differential CNV score* at bin *b* was defined as the difference between the mean alteration fraction of short PFI compared to medium and long PFI groups combined:

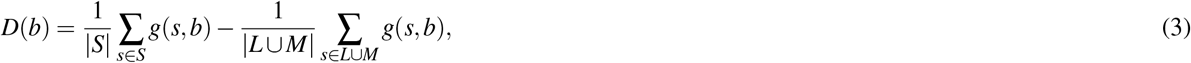

where *S* is the set of short PFI samples, and *L* and *M* are the sets of medium and long PFI samples. The score lies in [*−*1, 1], where positive values indicate alterations enriched in short PFI tumors.

A bin was flagged as short-PFI–enriched when it satisfied both of the following criteria: (i) at least *N* = 3 short-PFI samples showed an alteration fraction *≥*0.2 at that bin (recurrence criterion), and (ii) the differential CNV score exceeded 0.3 (differential criterion).

#### Minimal common region identification

To map candidate short PFI CNV enrichment loci to their minimal common region (MCR), flagged 1 Mb bins were re-examined at 100 kb resolution. For a given chromosome and CNV type, the per-sample gain (or loss) fraction was recomputed across 100 kb bins using the same subcluster-aggregation procedure described above. Contiguous flagged bins were merged into candidate blocks. For each block, the MCR was defined as the set of 100 kilobases (kb) bins satisfying two criteria: (i) the per-sample gain (or loss) fraction exceeded 0.1 in at least 3 of 4 short PFI patients, and (ii) the differential CNV score exceeded 0.2. Contiguous bins meeting both criteria were merged into a single MCR interval and reported with GRCh38 coordinates.

#### Tumor stroma clones

Analysis of the tumor-stroma border in the adnex cohort was performed with the stroma from all TMAs containing collectively as reference dataset, against all spots annotated as tumor stroma by the pathologist. InferCNV was first run with *HMM=FALSE*. From InferCNV’s hierarchical clustering (Ward’s D2) of the spots, visually distinctive clusters could be observed. The Visium spot barcodes that were part of such clusters were then extracted from the dendrogram, assigned a clone identity, and input to InferCNV again with *HMM=TRUE* to obtain high confidence CNVs from these clones.

### Kaplan-Meier analysis

The association between C3 expression and progression-free survival and overall survival in HGSC was analyzed using Kaplan-Meier Plot (https://kmplot.com/analysis/). The survival analysis was run using the two ovarian cancer datasets with the highest number of patients available, OV-TCGA^30^ and GSE9891^31^, both compiled of transcriptome gene-expression data. In the analysis, high C3 was defined using the “auto select best cutoff”, histology was set to “serous”, stage was set to “2,3 and 4” and grade to “2 and 3”. P-values were generated automatically by Kaplan-Meier using log-rank test. In both datasets, only treatment-naïve samples are included.

### Visium spot deconvolution

Cell types were mapped to the ST data using the cell2location^54^ (v0.1.4) Python package. To estimate the reference cell type signature, the gene expression data of a single cell V(D)J dataset from 14 ovarian cancer patients (223,363 cells) was leveraged^28^. The data were subset to cells from samples taken from the primary tumor and omental metastases, resulting in the exclusion of one patient, and a final cell count of 93,389, with 61,302 cells from primary tumor samples, and 32,087 cells from omental metastases. The reference data regression model was trained for 200 epochs with a batch size of 2,500. The final cell2location model was trained with hyperparameters *N_cells_per_location*=15, corresponding to the estimated mean number of cells per spot as assessed using QuPath (v0.5.1) cell detection, and *detection alpha*=20, for 3,000 epochs.

### RNA in-situ hybridization

The RNA-ISH was performed using the RNAscope 2.5 HD BROWN detection kit (#322300, Advanced Cell Diagnostics, Bio-Techne). In brief, 4 µm tissue sections were cut from PAXGENE fixed, paraffin-embedded TMA blocks, mounted on Superfrost slides, baked for 1h at 60 °C, and deparaffinized. After, the tissue sections were post-fixed with 4 % PFA, rinsed with sterile H2O and dried, followed by hydrogen peroxide treatment for 10 minutes at RT and 15 minutes target retrieval boiling at 98 °C. Then, tissue sections were treated with Protease Plus for 30 minutes at 40 °C and after, the target probes,C3 (#430701), IFI27 (#440111) and BST2 (#418653), were hybridized for 2 hours at 40 °C. The signal amplification steps were performed as described in the manual with the exception that Amp 6 was hybridized for 1 hour. Sections were treated with DAB for 10 minutes, counterstained with 50 % hematoxylin, dipped into ammonia water and coverslipped. The sections were scanned using the 3DHISTECH Panoramic 250 FLASH III digital scanner, a magnification of 40x was applied with extended focus and seven focus levels. Scanning was performed at Helsinki Biobank.

### RNA-ISH quantification

RNA-ISH signal was quantified using QuPath (v0.4.3). In short, every core was divided into quadrants and an 60 *µ*m times 60 *µ*m area of tumor epithelium from each quadrant was annotated. Further, mean DAB signal was quantified from each annotated areas using the QuPath Compute Intensity feature with a pixel size of 0.5 and a tile size of 15um. The average of the four quadrant scores (per core) was used for analysis; if patient had two cores from same tumor site, their average was used when plotting. Differences between PFI groups were determined by Kruskal-Wallis test followed by Dunn’s post-hoc test. Statistical analyses were performed in GraphPad prism (v10.6.1). An adjusted p-value < 0.05 was considered statistically significant.

## Author contributions statement

Conceptualization: N.A., A.R.L., O.M.C.; Methodology: F.N., N.A., S.H., O.Y., A.E., A.R.L.; Software: F.N.; Validation: F.N., N.A., A.E.; Formal analysis: F.N., N.A.; Investigation: F.N., A.E. ; Resources: N.A., S.H.; Data Curation: F.N.; Writing - Original Draft: F.N., N.A., O.Y., A.E.; Writing - Review & Editing: F.N., N.A., S.H., O.Y., O.M.C., A.E., A.R.L.; Visualization: F.N., N.A.; Supervision: O.M.C., A.E., A.R.L.; Project administration: A.E., O.M.C., A.R.L.; Funding acquisition: A.E., O.M.C.

## Acknowledgements

The spatial transcriptomics were performed at FIMM single-cell analytics and sequencing units supported by HiLIFE and Biocenter Finland. RNA quality control was provided by the Biomedicum Functional Genomics Unit at the Helsinki Institute of Life Science and Biocenter Finland at the University of Helsinki. RNA isolation, TMA construction and RNA-ISH slide scanning was performed at Helsinki Biobank which is supported by Hospital District of Helsinki and Uusimaa (HUS), University of Helsinki, Kymenlaakso Social and Health Services (Carea) and the South Karelia Social and Health Care District (Eksote). Funding was received from the Research Council of Finland (A.E. 360763), the European Union Horizon 2020 research and innovation program (O.M.C. Grant Agreement No 965193 for DECIDER), Instrumentarium Tiedesäätiö (A.E. 240003), Sigrid Juselius Foundation (A.E. 250031 and O.M.C.), Cancer Foundation Finland (A.E. 63-6403 and O.M.C.), Finska läkaresällskapet (O.M.C.). F.N. received personal funding from the iCANDOC national doctoral education pilot in precision cancer medicine, Finland. N.A. received personal funding from the Faculty of Medicine, Doctoral Program in Clinical Research. A.L. salary was partially supported by the iCANPOD postdoctoral program, which is funded through the iCANDOC doctoral education pilot in precision cancer medicine. O.Y. received personal funding from K. Albin Johanssons Stiftelse.

## Additional information

Supplementary Information is available for this paper.

## Data availability

The data generated and analyzed in this study were deposited in Zenodo and are available at the following URL: 10.5281/zen-odo.22746547. During peer review, the files are held under restricted access and are available to editors and reviewers via a confidential access link; they will be made publicly available upon publication.

## Code availability

The code for all the analyses in this paper are available at https://github.com/Franzili/spatial-hgsc-interferon.

