## Supplementary for "Spatial transcriptomics reveals interferon pathway enrichment in early relapsing serous carcinoma"

### Supplementary figures

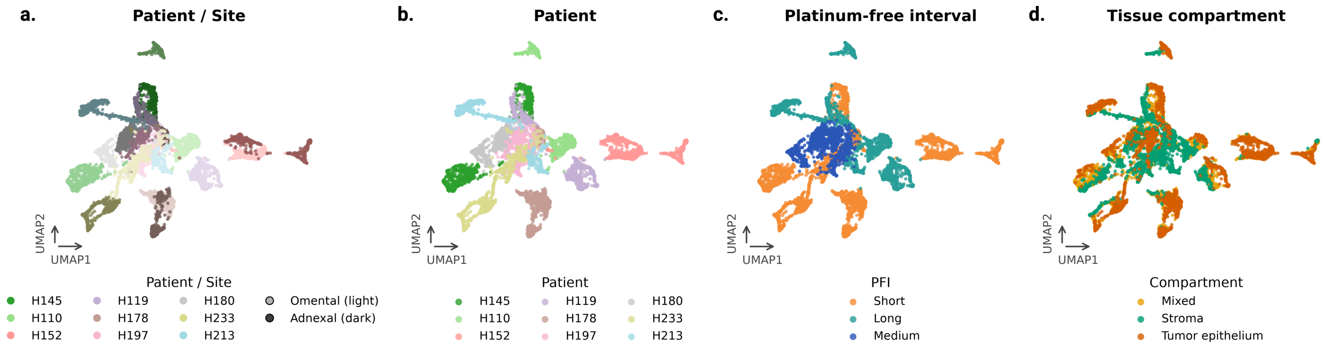


**Figure 1. UMAP projections.** UMAPs showing the distribution (from left to right) of patients and tumor site (**a.**), patients (**b.**), PFI (**c.**), and tissue compartment (**d.**). PFI, platinum-free interval; UMAP, Uniform manifold approximation and projection.

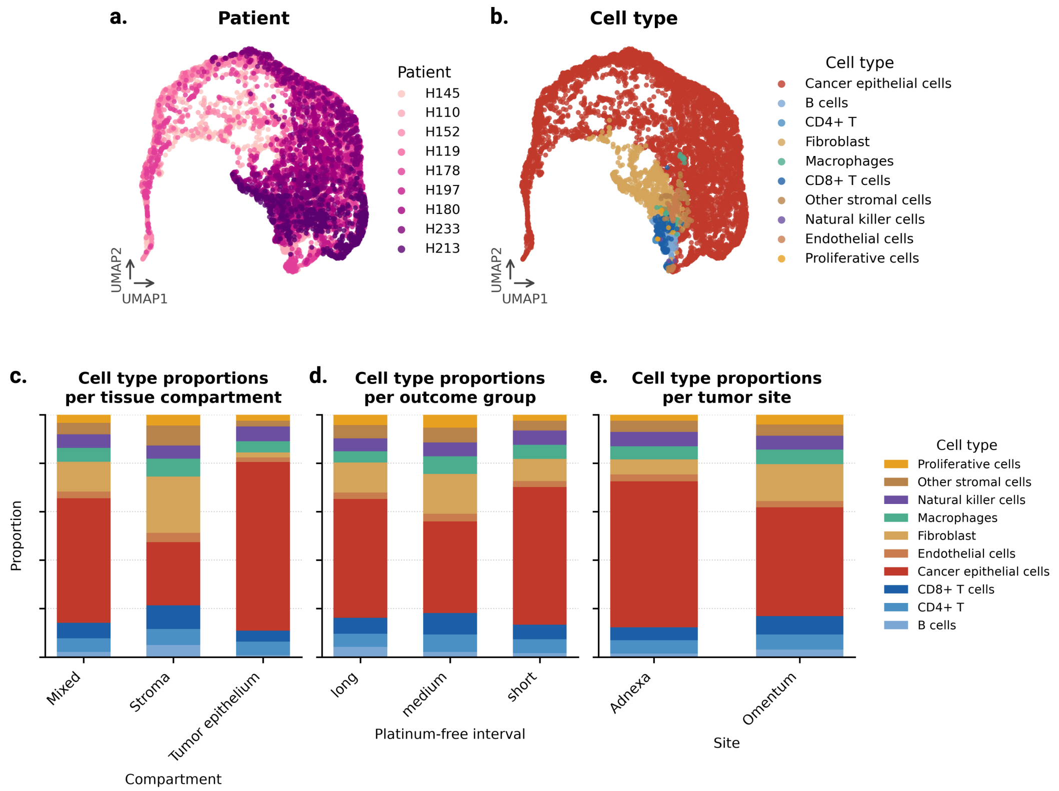


**Figure 2.** **Cell type deconvolution.** UMAP projections showing the distribution of patients (**a.**) and inferred cell types (**b.**). Cell type proportions vary by tissue compartment (**c.**), patient outcome (**d.**) and tumor site (**e.**). UMAP, Uniform manifold approximation and projection.

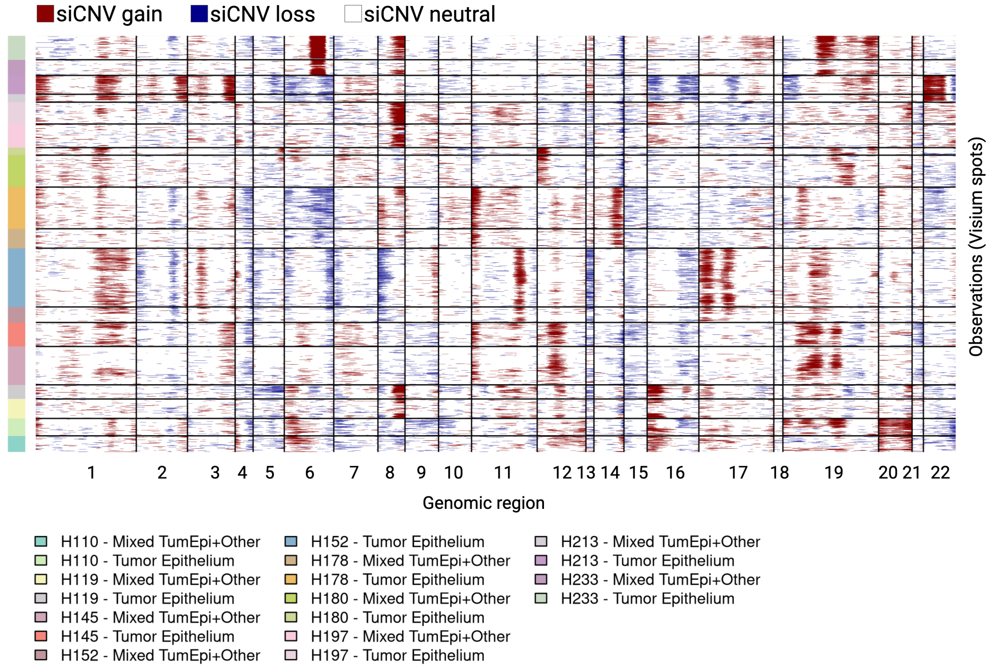


**Figure 3.** **Copy-number profiles of the tumor epithelium.** Spatially inferred copy-number variations (siCNVs) from tumor epithelium and mixed Visium spots of all patients. As reference, stromal spots from all patients collectively have been used.

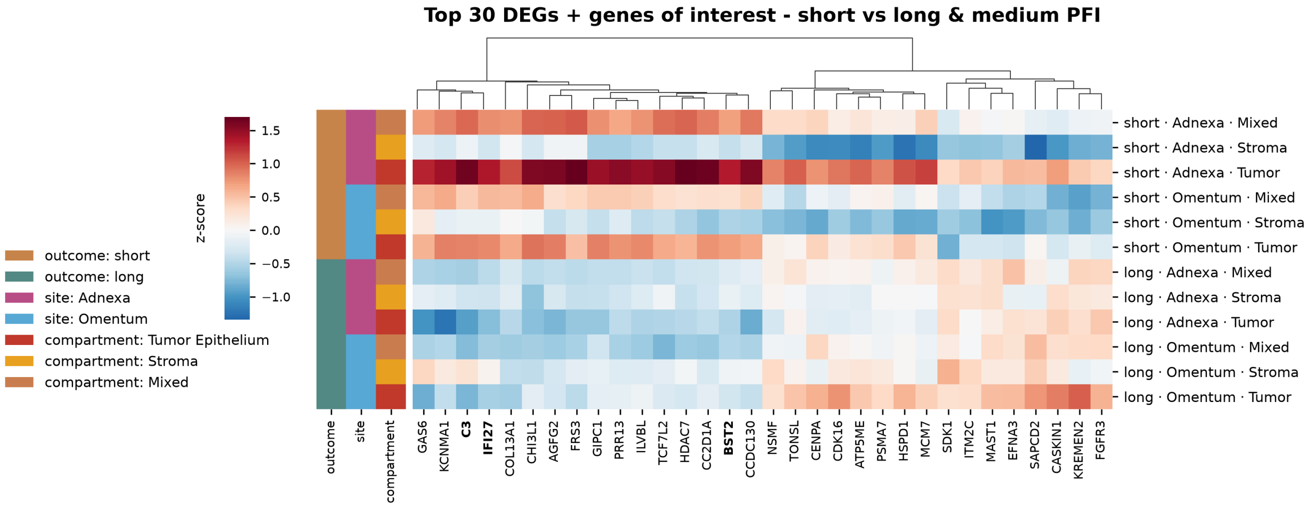


**Figure 4.** Top 30 differentially expressed genes in the short PFI group as compared to the med/long PFI group. Bolded gene names: genes of interest selected for deeper analysis. PFI; platinum free interval

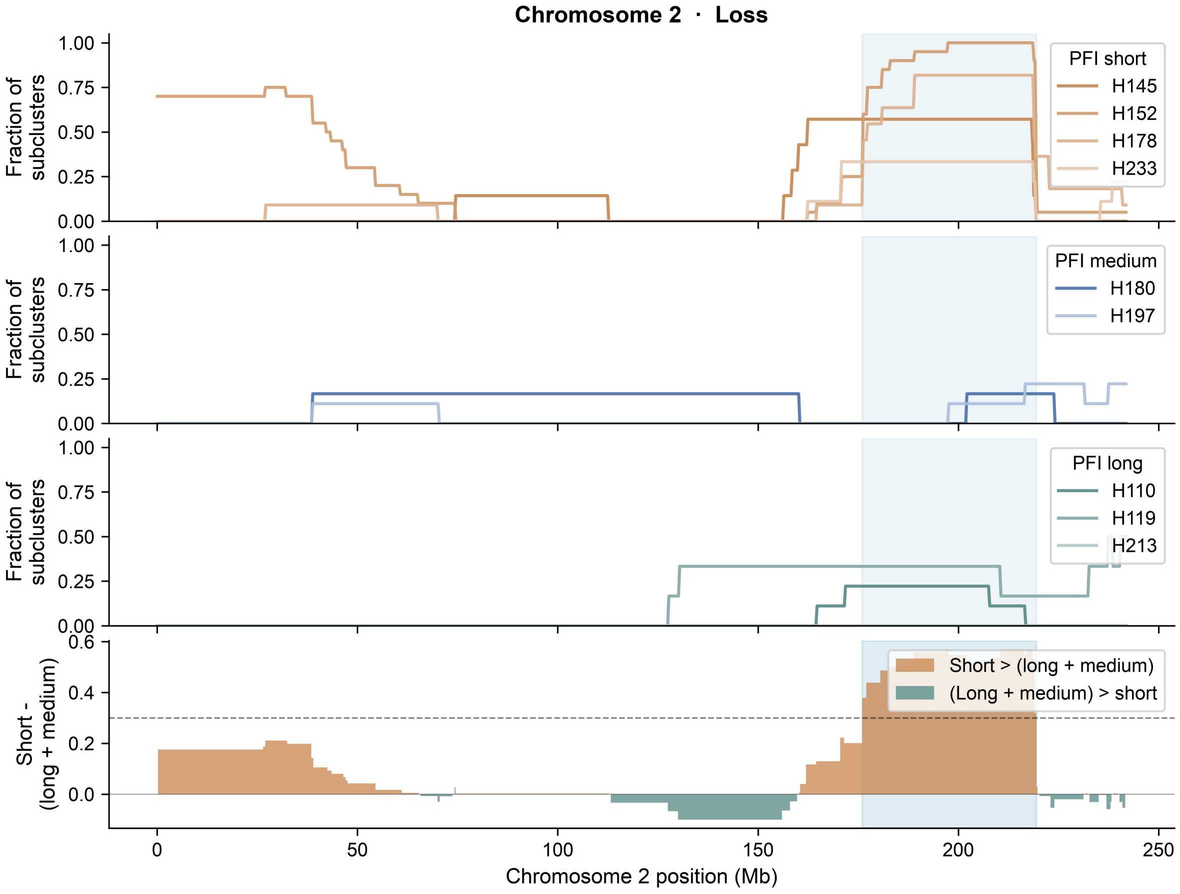


**Figure 5. CNV profile for Chromosome 2 losses enriched in short PFI patients.** Per-patient loss fraction profiles are shown at 100 kb resolution. Each line represents one patient, colored in shades of the corresponding PFI category (short: orange; medium: blue; long: green). The lowest panel shows the differential CNV score (mean short PFI minus mean (long+medium)-PFI loss fraction) at 1 Mb resolution; orange fill indicates regions where short PFI exceeds (long+medium)-PFI, green fill the reverse. Shaded vertical bands denote candidate minimal common regions (MCR). Vertical dotted lines indicate the positions of genes of interest.

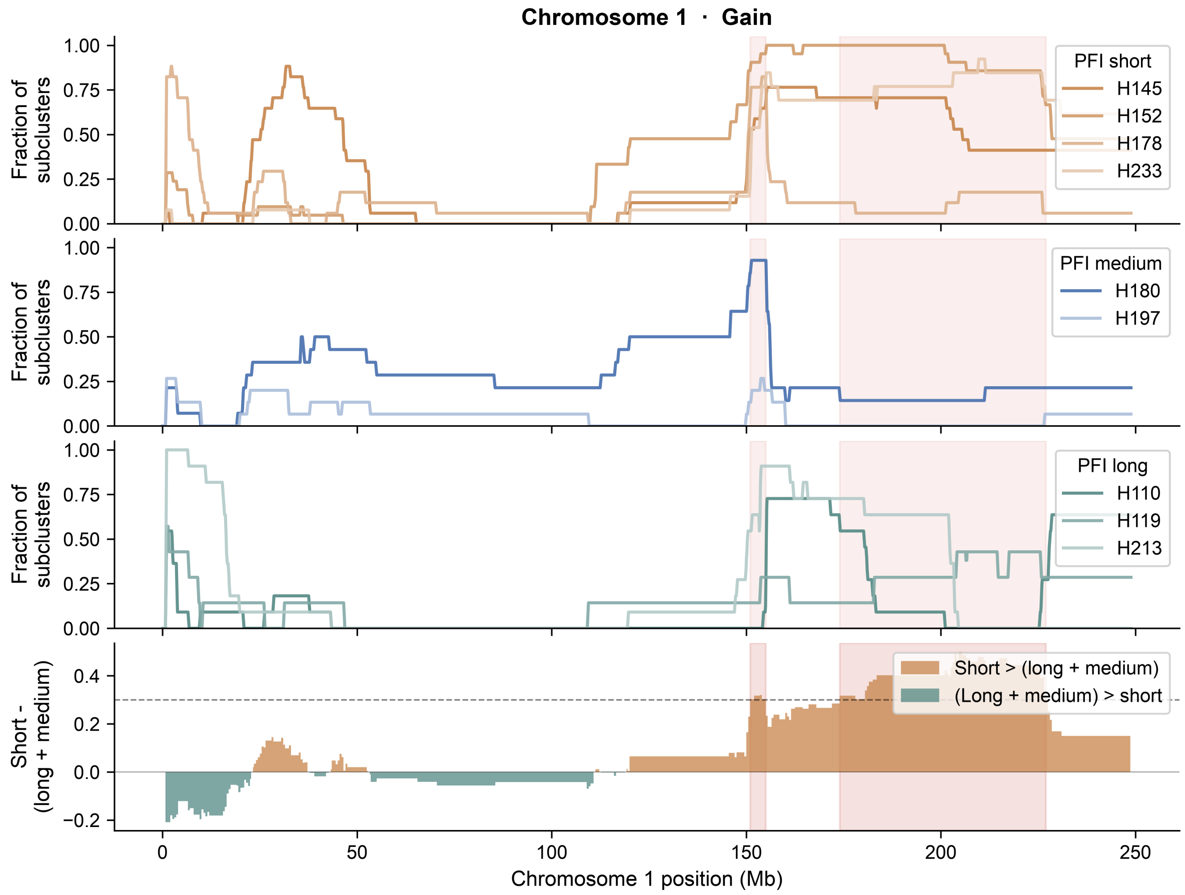


**Figure 6. CNV profile for Chromosome 1 gains enriched in short PFI patients.** Per-patient loss fraction profiles are shown at 100 kb resolution. Each line represents one patient, colored in shades of the corresponding PFI category (short: orange; medium: blue; long: green). The lowest panel shows the differential CNV score (mean short PFI minus mean (long+medium) PFI loss fraction) at 1 Mb resolution; orange fill indicates regions where short PFI exceeds (long+medium)-PFI, green fill the reverse. Shaded vertical bands denote candidate minimal common regions (MCR). Vertical dotted lines indicate the positions of genes of interest.

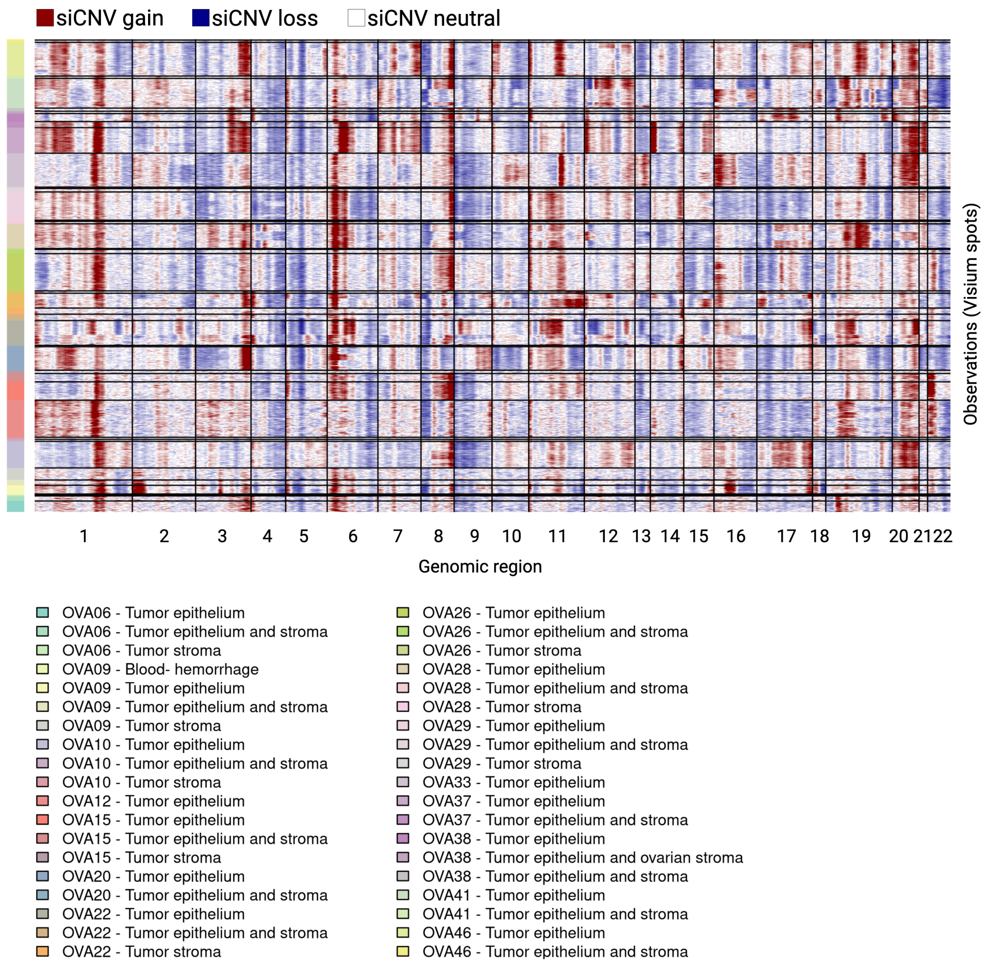


**Figure 7. Copy-number profiles of an external HGSC cohort.** Spatially inferred copy-number variations (siCNVs) from Visium spots annotated by a pathologist as tumor epithelium, tumor epithelium and stroma, tumor stroma, and blood-hemorrhage. Spots annotated as ovarian stroma from all patients collectively were used as reference for inference.

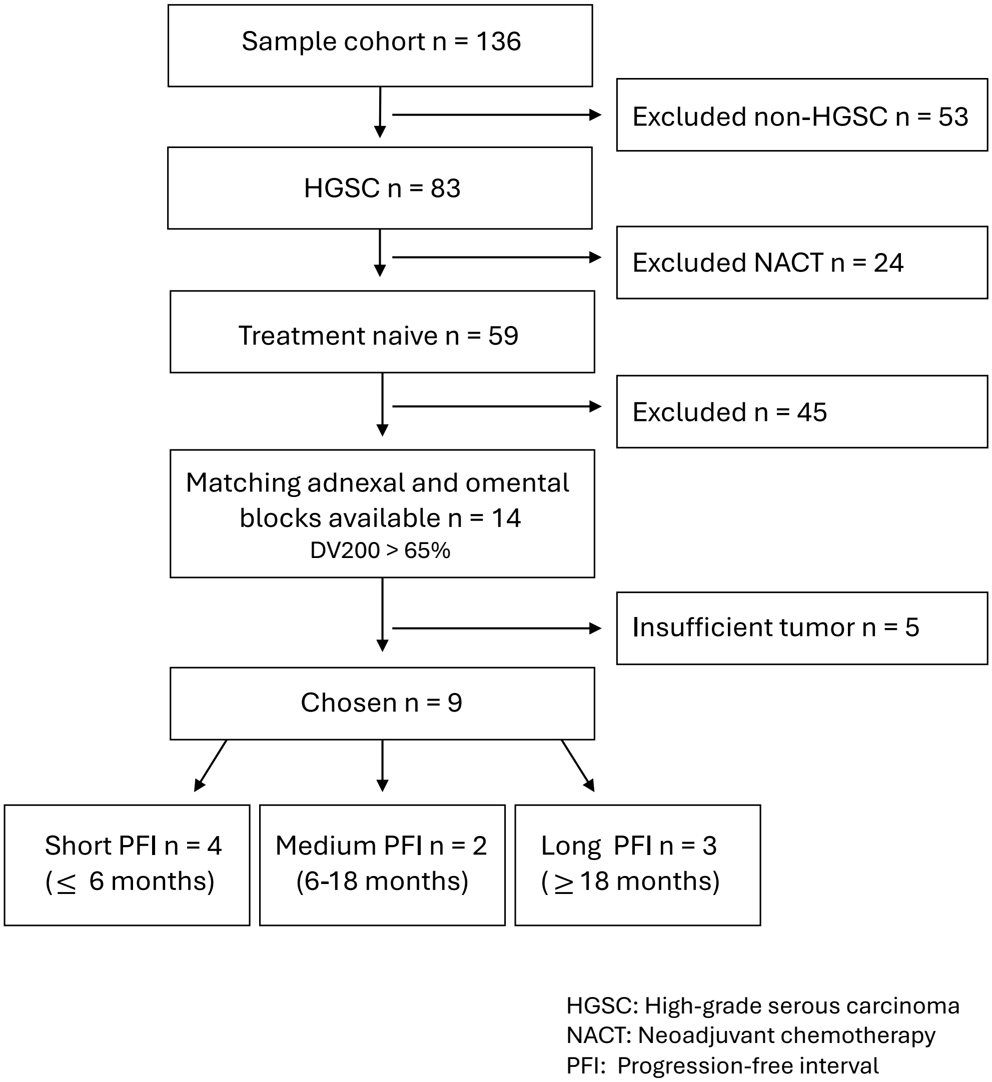


**Figure 8.** **Full cohort filtering criteria.**

### Supplementary Results

Of the 13 antigen-presentation genes in the Visium probe set that were measured in our data, only CD74, also known as HLA-DR, appeared in the leading edge (7.7%), compared with 51 of 187 remaining set members (27.3%). No class I or peptide-loading gene reached the leading edge in either interferon set (0 of 8 for interferon gamma, 0 of 5 for interferon alpha), and four of the five measured class II genes (CIITA, HLA-DMA, HLA-DQA1 and HLA-DRB1) were likewise absent. Testing depletion of the antigen-presentation module against the remaining set members gave a one-sided Fisher $p = 0.021$ (odds ratio 0.13) for interferon gamma. The interferon alpha set contains only six measured antigen presentation genes and could not support the test ($p = 0.112$).

As an independent check at the level of whole gene sets, we ranked all Hallmark sets by class II content. The most class II–rich set, allograft rejection (7 of 14 measured class II genes), was not enriched between outcome groups (NES = 1.00, nominal $p = 0.45$, FDR $q = 0.76$), despite the strong enrichment of both interferon sets in the same comparison. Notably, four class I genes (HLA-A, HLA-E, HLA-G, TAPBP) did reach the allograft rejection leading edge, indicating that the exclusion we observe is specific to the interferon program rather than a global property of antigen-presentation genes.
